# MIMIQ: Fast mutual information calculation and significance testing for single-cell RNA sequencing analysis

**DOI:** 10.64898/2026.04.10.717770

**Authors:** Daniel O’Hanlon, Sergio Garcia Busto, Rubén Pérez Carrasco

## Abstract

Mutual information is a fundamental quantity in information theory that describes the non-linear dependency between two variables, and has numerous applications within bioinformatics and beyond. However, its exploitation is hampered by a trade-off between computational intensity and accuracy. Here we present an adaptive binning approach to computing the pairwise mutual information, optimized for small integer counts such as those observed in single-cell RNA sequencing. By assuming a sampling distribution such as the negative binomial, a χ^2^ test statistic for hypothesis testing can be computed simultaneously via a copula transformation. Using these quantities, we show how gene rewiring of CD4+ naive T-cells during SARS-CoV-2 infection can be studied using a single-cell sequencing dataset of healthy and COVID-19 donors.

## 1 Introduction

The calculation of gene covariation across sequenced cells is an essential component of numerous analytical pipelines, from cell type inference and clustering to regulatory network and trajectory inference [1–3]. The most straightforward metrics of this covariation are the Pearson or Spearman correlation coefficients. However, these are unable to identify complex non-linear dependencies that are characteristic of multiple interposed genetic interactions. The mutual information (MI) between a pair of variables describes the information content gained about one variable when measuring the other, and is a model-independent metric of the interaction. Furthermore, MI itself is a meaningful quantity, particularly when investigating information flow in signal transduction [4, 5], or the diversity of immune receptors [6].

Precise calculation of the MI between all pairs of genes is computationally intensive, and is therefore unsuitable for applications involving the tens of thousands of genes (hundreds of millions of gene pairs) detected in a modern single-cell RNA sequencing (scRNA-seq) experiment. Conversely, faster but more approximate methods that calculate the MI in fixed bins often lack accuracy when distributions are skewed, such as those in RNA sequencing, which are predominantly described by a long-tailed overdispersed distribution.

Here we present MI from Marginally Informed Quantities (MIMIQ), a framework for the fast calculation of pairwise MI, without compromising accuracy, and with light assumptions on the underlying data distributions. The approach is optimized for use with potentially zero-inflated integer count data, such as those obtained in scRNA-seq experiments, which can be described via zero-inflated negative-binomial distributions. To achieve this, we use a k-d tree to perform adaptive binning of the raw counts, combined with a copula transform that enables simultaneous calculation of a *χ*^2^ test statistic for significance testing. To illustrate the usage of these quantities in practice, we formulate an MI-based rewiring metric and use it to investigate gene interaction rewiring of naive T-cells during SARS-CoV-2 infection.

The algorithm described in this paper is available in C++ with a Python interface at github.com/dpohanlon/mimiq, and as the Python package `mimiq` on PyPI.

## 2 Previous work

One of the fastest ways to estimate discrete pairwise MI is the naive ‘plug-in’ maximum-likelihood estimator, which computes the empirical joint and marginal probabilities and substitutes them directly into the MI formula [7, 8]. Its accuracy depends on how the contingency table is populated: when the effective support of the joint distribution is large relative to the sample size, many joint states are rare or unobserved, leading to high variance and finite-sample bias. Coarsening the data by uniform binning can reduce variance, but introduces resolution-dependent bias and can behave poorly for heavy-tailed or highly skewed count distributions such as the (zero-inflated) negative binomial. Conversely, k-nearest-neighbour (kNN) estimators [9, 10] infer MI from local density ratios and are broadly applicable and consistent for continuous data, but are substantially more expensive to compute.

To achieve a balance between computational efficiency and modelling accuracy, we employ a copula framework that separates marginal distributions of each variable from dependence between variables [11]. This approach constructs a joint distribution by first establishing a distribution with uniform marginals that capture the dependence structure of the data. These uniform marginals are then transformed into the desired target distributions using their known inverse cumulative distribution functions (CDFs). Here we pair a non-parametric joint distribution constructed from k-d trees [12] with zero-inflated (ZI) negative-binomial (NB) marginal distributions. The ZINB distribution is a mixture of a negative-binomial component, which captures biological variability [13], and a point mass at zero, which represents technical dropouts. By modelling dropouts explicitly, we can better isolate and characterize the underlying biological count distribution, although this is likely unnecessary for counts based on unique molecular identifiers [14].

For each gene *g*, we calculate an estimate of the marginal CDF, either by fitting one of these parametric distributions to the data, or by estimating the CDF empirically. Such an estimate is preferred when the fit quality of the parametric distribution is poor. For each gene *g*, we construct an empirical marginal CDF from the observed integer counts. We first tabulate the number of observations at each count level, then add a small pseudocount *α* (here *α* = 0.5) for smoothing. After normalisation, the empirical CDF is obtained by cumulative summation,

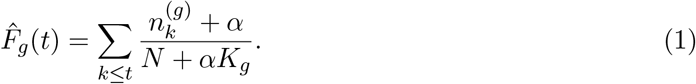

Here, *N* is the total number of observations, 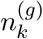 is the number of observations for which gene *g* has count exactly *k*, and *K*_*g*_ is the number of integer count levels in the observed range of gene *g*.

## 3 Pairwise MI calculation

For a gene pair (*X*_*a*_, *X*_*b*_), we build a balanced k-d tree in the original (*X*_*a*_, *X*_*b*_) count space by alternating median splits until no leaf contains more than *N*_leaf_ unique observations. An example of this binning for simulated data can be seen in Figure 1. The resulting leaves 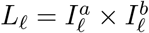 for *ℓ* = 1, …, *L*, tessellate the joint count space into rectangular regions defined by the gene-specific intervals *I* = [*I*^−^, *I*^+^]. The empirical joint probability for an observation in leaf *ℓ* is given by 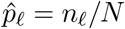 where *n*_*ℓ*_ is the total number of observations in leaf *ℓ*. Similarly, we can calculate the marginal distribution for each interval *I*, using the estimated marginal CDF 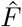 for each gene over the corresponding bin interval *I* = [*I*^−^, *I*^+^]. For instance, for gene *a* we have

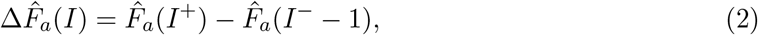

and similarly 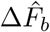. The ‘area’ of this leaf is therefore

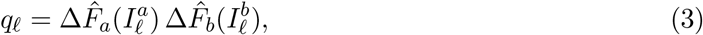

which corresponds to the product of the marginal probabilities. With these probability estimates, we can use the plug-in MI estimator in each bin to calculate the total pairwise MI,

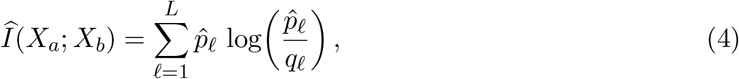

which describes how much the bin contents deviate from that expected from the product of marginals.

**Figure 1:**
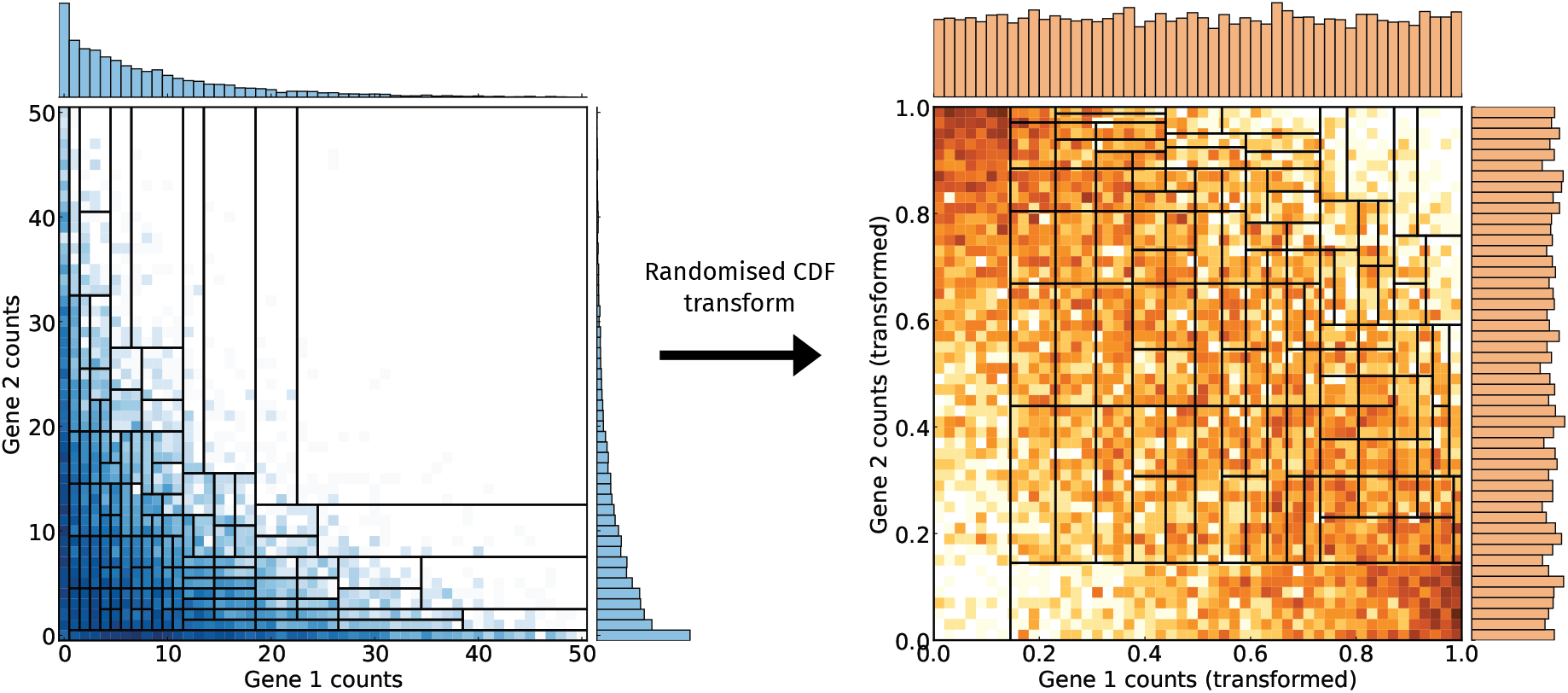
The k-d tree binning procedure applied to a simulated dataset with 5% zero-inflated NB marginal distributions, and a joint normal distribution with −0.5 correlation. Darker colours indicate more populated bins, and the shape of each marginal distribution is located opposite the axis labels for that axis. Left) histogram of the raw data and adaptive binning (black lines) in count space. Right) Histogram of the data and binning after the randomized CDF transformation according to the NB marginals. After the transformation, the marginals are uniformly distributed, and the bin width along each axis is proportional to the probability mass within that bin according to the NB distribution.

One advantage of this construction is that after the CDF transformation we expect independent pairs (i.e., with *I*(*X*_*a*_; *X*_*b*_) = 0) to be uniformly distributed, so we can construct a *χ*^2^ test-statistic as

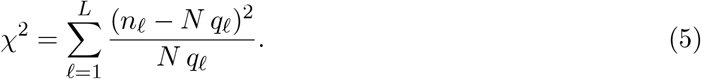

### 3.1 Hypothesis testing

To estimate the test statistic, we randomly split the dataset into two sets. The first set is used to construct the adaptive binning, while the second set is used to evaluate the *χ*^2^ statistic. Under the null hypothesis, the binning scheme is independent of the value of the test statistic, and therefore the expected distribution is a *χ*^2^ distribution with a number of degrees-of-freedom equal to *L* − 1. This circumvents the need to estimate the effect of the adaptive binning on the effective degrees-of-freedom. Much like in *k*-fold cross validation, we can also reverse the roles of the two subsets, by constructing the binning based on the second subset and evaluating the statistic on the first. The resulting *χ*^2^ values and their degrees of freedom can then be summed to obtain a combined test statistic.

To evaluate the performance of the method, we analyse the probability of incorrectly rejecting the zero MI null hypothesis (type I error). Figure 2A shows the type I error for a p-value of 0.05. The results are obtained for ensembles of pairs of independent NB marginal distributions. As expected, the type I error is clustered around the expected *p* = 0.05 value independently of the number of observations.

**Figure 2:**
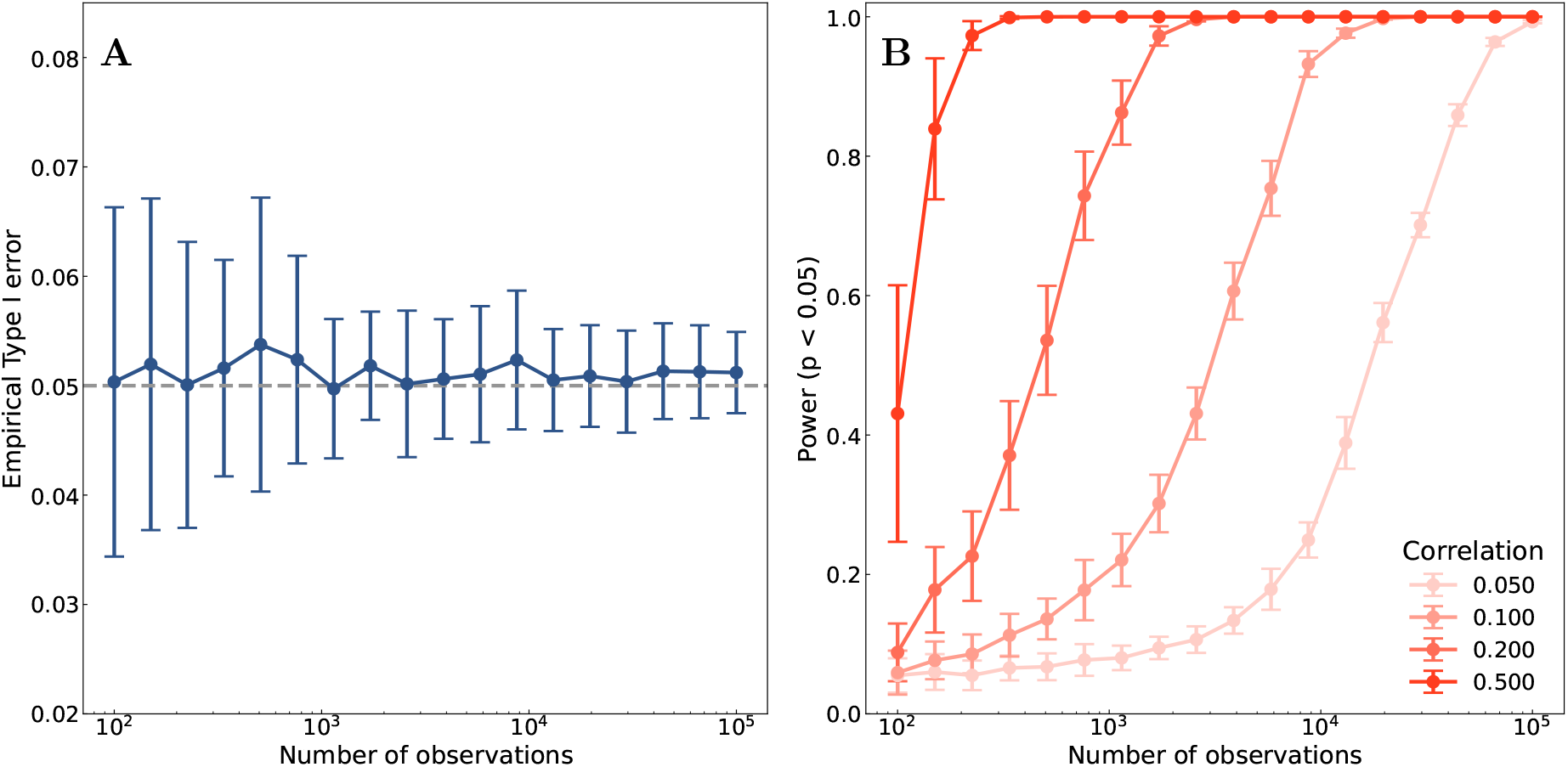
Hypothesis test type I error (**A**) (with a p-value of 0.05) and test power (**B**) for the data with NB marginal distributions, as a function of the number of observations. The type I error is reasonably constant with the number of observations, and distributed around the desired p-value of 0.05. This data includes features with known correlation in order to estimate the increase in power when the correlation is increased, and with an increasing number of observations. In both panels, 50 *×* 50 pairs of NB distributions are generated by uniformly sampling *r* ∈ [1, 20] and *p* ∈ [0.05, 0.7] with a minimum leaf count of 50.

Furthermore, the converse probability of correctly rejecting the zero MI null hypothesis (the test power) is shown in Figure 2B for various generated values of the correlation between the NB distributions. The results show that the power increases with both the correlation parameter of the joint distribution and the number of observations.

## 4 Benchmarks

Due to the non-linear form of the MI, closed-form expressions are generally unavailable except for a few special cases. To assess the accuracy of our estimator we therefore consider the case in which pairs of variables are coupled through a Gaussian copula with correlation parameter *ρ*, whose marginals are subsequently zero-inflated. In this setting the MI can be approximated (Appendix A) as

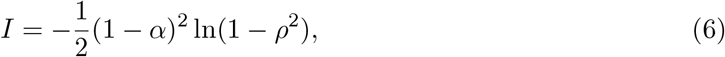

where *α* denotes the zero-inflation of each variable.

We compare MIMIQ with the kNN implementation in scikit-learn [9, 10], and a maximum-likelihood estimator designed for large-scale gene expression datasets with a Miller-Madow bias-reduction correction [7, 15] (FastGeneMI). We compare the accuracy of these methods with respect to the analytical expectation (6), using distributions that are correlated through a Gaussian copula for differing marginal distributions: negative-binomial, and zero-inflated negative-binomial. For the MIMIQ method, the minimum bin content is set to be 50. Synthetic data were generated by constructing a set of 50 interdependent features, with dependence induced by a covariance matrix of rank 10. Specifically, samples were drawn from a multivariate normal distribution with covariance written in factorized form, Σ = *LL*^⊤^ + *D*, where *L* is a 50 *×* 10 loading matrix and *D* is a diagonal noise term. The resulting data were then transformed to impose independent marginal distributions while preserving the Gaussian copula. MIMIQ achieved an accuracy comparable to the scikit-learn kNN method, while improving on the FastGeneMI estimates (Figure 3A). The deviation from the analytical prediction decreases as the number of observations increases independently of the marginal distributions of the data, converging to the analytical expectation (Figure 3B).

**Figure 3:**
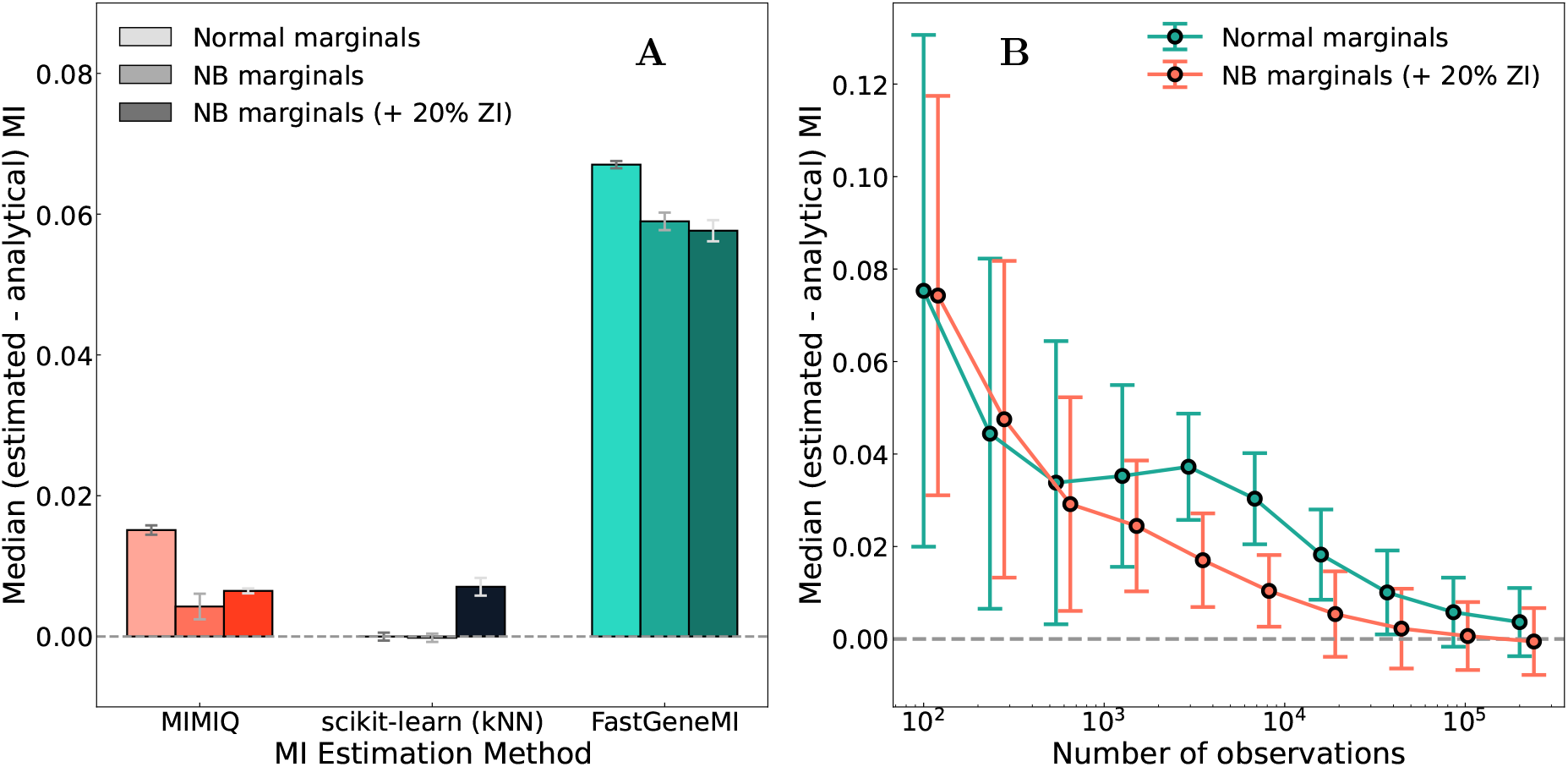
Comparison between analytical and estimated MI (Eq. (6)). In **A**, the MI error obtained using different methods with respect to the analytical prediction for a selection of marginal distributions. The correlation is generated by Gaussian copula with 10^4^ observations, and both the scikit-learn and FastGeneMI functions are run using their default arguments. The accuracy of MIMIQ (**B**) increases with the number of observations, converging to the analytical value independently of the marginal distributions. Error bars indicate standard deviations.

In addition to the accuracy we also analysed the runtime performance for the different pairwise MI algorithms (Figure 4). This was executed on an 11 core M3 MacBook Pro, with 18GB of RAM and the number of OpenMP [16] threads `OMP_NUM_THREADS` set to 8. As expected, the kNN approach is significantly slower (two orders of magnitude) than either MIMIQ or the naive plug-in estimator approach, FastGeneMI. Whilst all approaches scale according to the number of gene pairs, 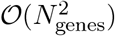, the kNN algorithm involves more computations per pair, as it calculates local density estimates after coarsely partitioning the space.

**Figure 4:**
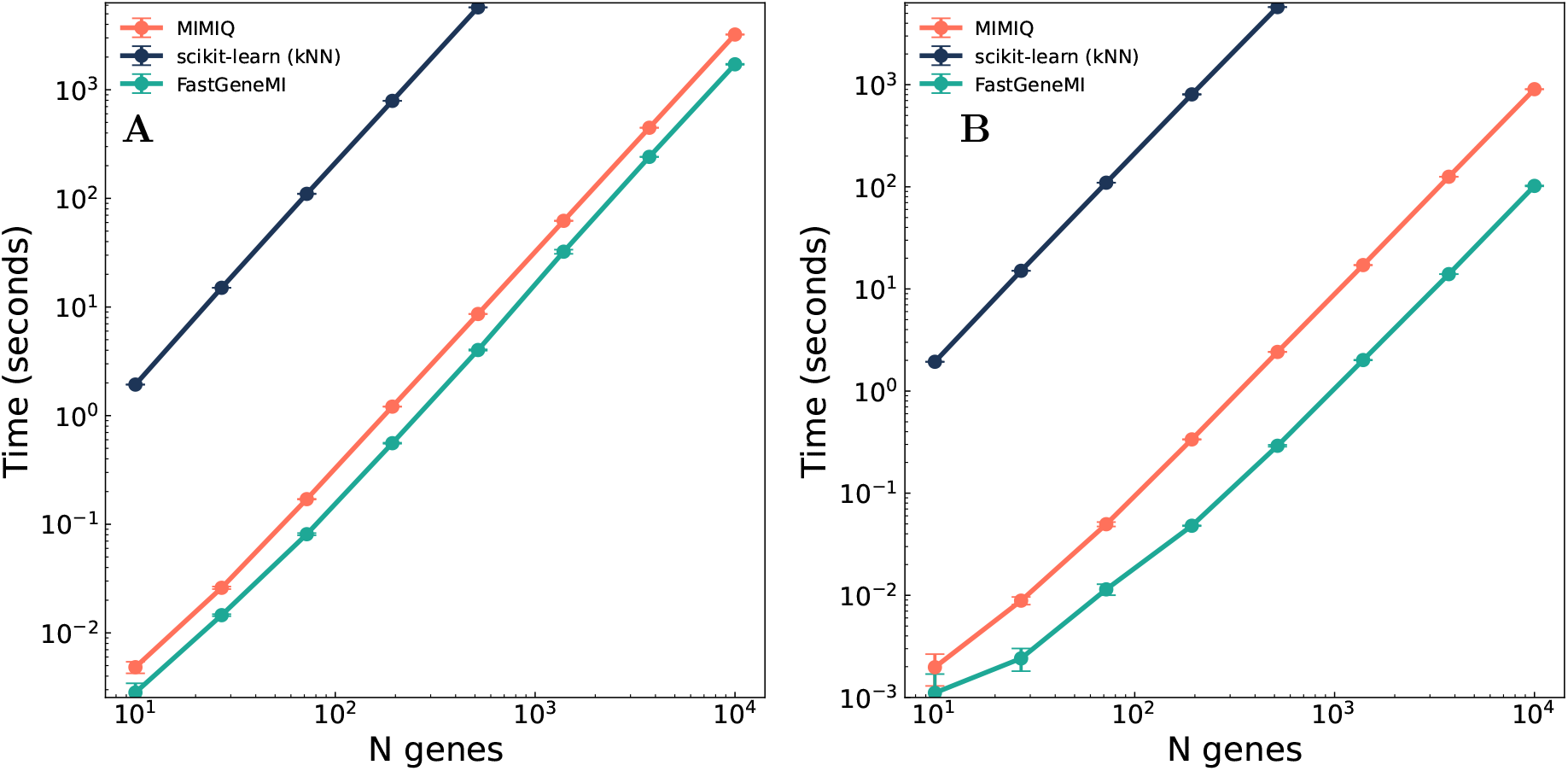
The run times of the algorithms presented as a function of the total number of unique features for 10, 000 observations, with the same sampling distributions as those in Figure 3. This is shown separately for normally distributed marginals (**A**) and negative-binomial marginals (**B**), where the central values are the median of 3 runs and the uncertainties are the standard-deviation of these.

## 5 Application to single-cell sequencing data

To evaluate the performance of the algorithm on real data, we calculate the MI between highly variable genes expressed in the Yoshida *et al*., peripheral blood mononuclear cell (PBMC) dataset [17]. This dataset comprises single cell RNA sequencing of PBMCs for adult and paediatric participants infected with SARS-CoV-2, along with corresponding healthy controls.

In total, the dataset comprises approximately 422, 000 cells across cell types and conditions. We apply a standard scRNA-Seq pre-processing pipeline [3, 18] to remove low-quality cells and reject droplet doublets with Scrublet [19]. This preserves only those cells with greater than 200 counts, only those genes with counts present in three or more cells, and removes cells that have in excess of 20% of their transcripts corresponding to mitochondrial genes.

To establish the accuracy of MIMIQ on real data, we start by assuming that the scikit-learn kNN estimator is an accurate representation of the MI in this data. This is applied to the 2, 000 most highly variable gene pairs within the SARS-CoV-2 dataset, selected after normalization and transformation of the counts, followed by highly-variable gene selection [3]. This is compared with the output of MIMIQ, where both methods are evaluated on the raw transcript counts. For MIMIQ, the marginal NB distributions are fitted using gradient descent, assuming zero inflation and where the parameters are initialized using the sample mean and variance. The maximum bin occupancy for the k-d tree is set to 50. For this comparison, both the scikit-learn kNN estimator and MIMIQ are evaluated on the raw unnormalized and untransformed integer transcript counts, for those genes and cells that pass the above pre-processing pipeline, to ensure consistency with the NB assumptions.

We investigate how the agreement between both methods varies as a function of the MI between the transcript counts of gene pairs. To this end, gene pairs are grouped into 8 bins according to their MI. The difference between both MI estimates is displayed in Figure 5A. Good agreement is observed across the range of MI observed in this dataset, [0.0, 0.43].

**Figure 5:**
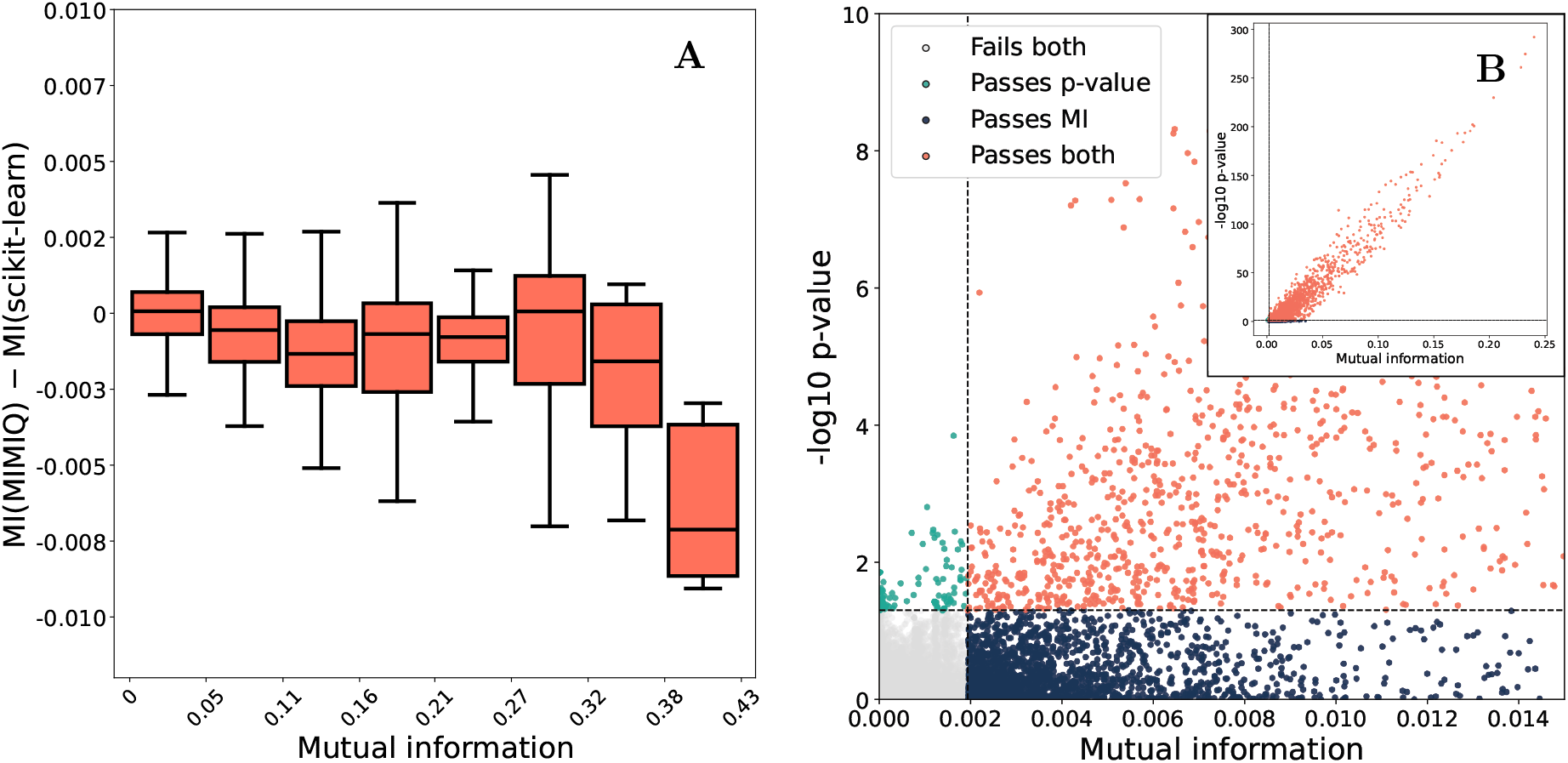
In **A**, box plots indicating the distribution of differences between the scikit-learn kNN MI estimator and the MIMIQ estimator, in units of MI bits. The thick bar inside the box indicates the median value of this difference, the box indicates the interquartile range (IQR), and the whiskers extend to 1.5×IQR. In **B**, for COVID-19 positive samples, the MI versus the negative logarithm of the p-value for the pairs of genes selected by the pre-processing selection. The selection region used for the subsequent rewiring analysis is indicated, and the inset is the full distribution. Pairs retained for subsequent analysis are shown in red, whereas rejected pairs are shown in grey, teal, and dark blue. Approximately 50% of these pairs that would otherwise pass the requirement on the MI fail the requirement on the p-value (dark blue) and are subsequently rejected.

To exemplify how MI calculations utilized across large datasets, we investigate rewiring of gene relationships in CD4+ naive T cells, under two conditions: for healthy and COVID-19 positive donor cells. Before the QC and pre-processing steps, there are 31, 109 COVID-19 donor cells and 46, 330 non-COVID-19 donor cells (including those identified as ‘post’ COVID-19). These are cells identified as naive CD4+ T cells by the cell-type labelling in the original analysis [17], and to these we apply the aforementioned QC pipeline. To purify the CD4+ T-cell selection, we reject cells consistent with erythrocyte markers (such as expression of the haemoglobin subunits *HBA1/2* and *HBB*), and those with B-cell markers (such as *CD79A*) and immunoglobulin heavy and light chain genes. We further remove housekeeping, ribosomal, and mitochondrial gene transcripts, from the remainder select only 2, 000 highly-variable genes for further analysis. To exploit the properties of the negative-binomial distribution and perform the MI estimation on raw counts, yet still correct for library size differences between cells, we perform binomial thinning of the data. Cells with transcript counts considerably larger than the median have transcripts dropped randomly, such that the total count is comparable across cells.

The advantage of having access to both the MI and the statistical significance is that pairs with high MI but low significance can be rejected, resulting in a more robust analysis. The rejection of these pairs can be visualized in Figure 5B, where the requirements for both MI and statistical significance result in 50% fewer spurious associations than a requirement for MI alone.

For each pair of conditions, *C* ∈ {*C*_0_, *C*_1_}, and for a gene *g*_*i*_, we compute a rewiring score, *R*_*i*_, over all other genes *g*_*j*≠*i*_ with significant MI (*p <* 0.05) under both conditions. This is the sum of the absolute difference of the conditional MI values between the two conditions,

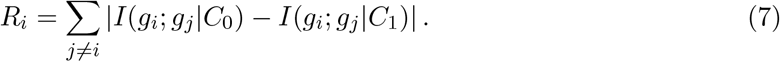

This quantifies the extent to which the local interaction environment of *g*_*i*_ changes between conditions. Between COVID-19 and healthy donor cells in naive CD4+ T cells, we find that *ZFP36*, a negative feedback regulator of T-cell response is the most rewired gene (Figure 6). The rest of the top rewired genes are dominated by established T-cell activation markers, such as *FOS, JUN, TNFAIP3*, and *NFKBIA* [20–22]. The highest MI interactors with *ZFP36* are dominated by immune signalling genes [23, 24]. When comparing the pattern of interactions within the ego network of *ZFP36*, higher MI is observed particularly between *ZFP36, NFKBIA*, and *DUSP1*, key regulators of immune response signalling [25]. This signalling and regulation scheme is recapitulated in the complementary differentially expressed gene analysis, where many key controllers of T-cell activation are upregulated in COVID-19, and the proliferation regulator *TXNIP* is downregulated [26].

**Figure 6:**
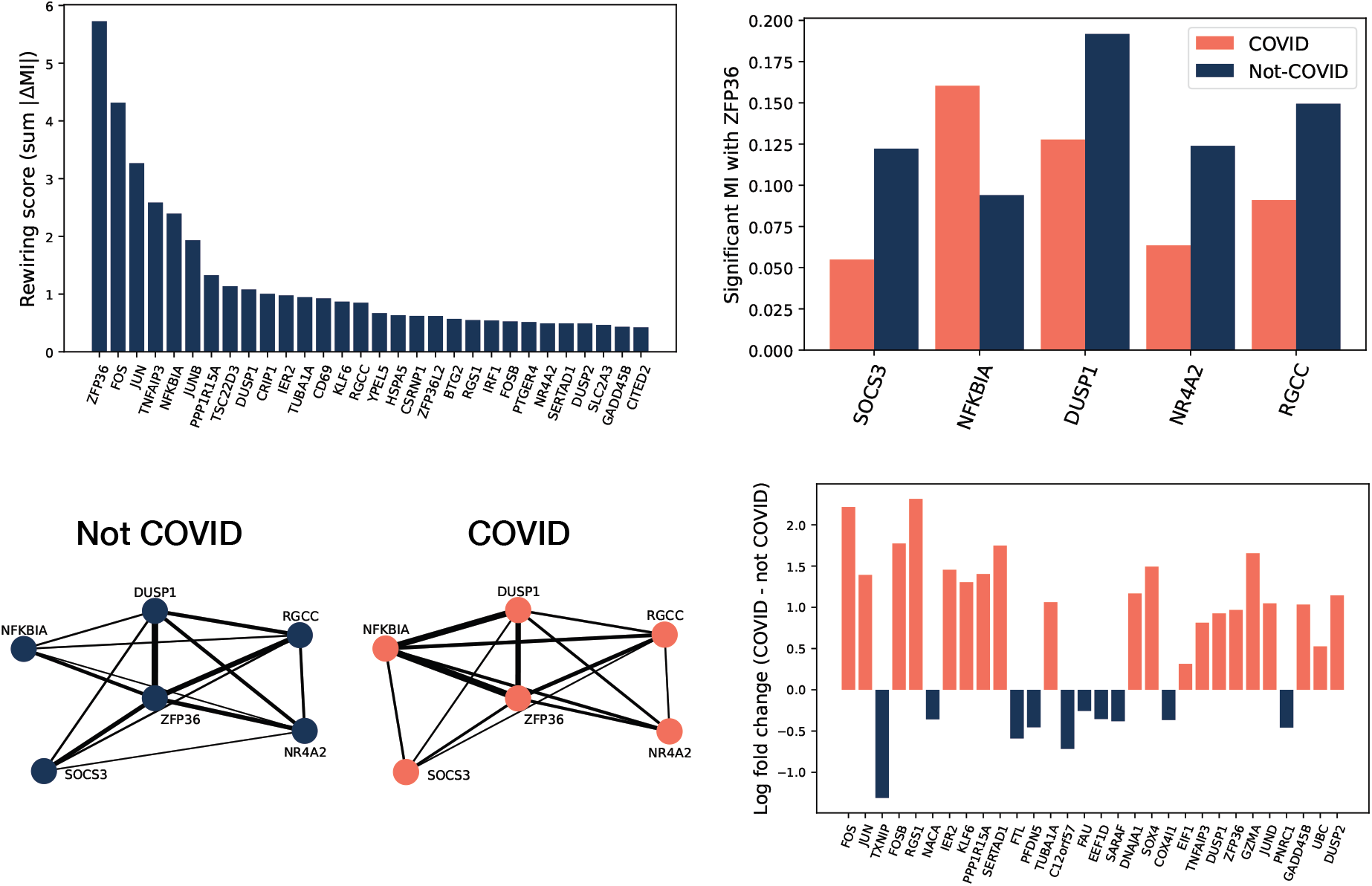
Gene rewiring for naive CD4+ T cells, between cells from donors in the COVID-19 versus healthy cohorts. The top rewired genes (top left) according to the rewiring score are dominated by T-cell activation markers. The highest MI interactors with the most rewired gene according to the rewiring score (top right), *ZFP36*, are dominated by immune signalling genes. The difference between the interactions of these genes in the COVID-19 and non-COVID-19 cohorts is displayed graphically (bottom left), where edge thickness is proportional to the MI. In the COVID-19 cells, more interaction is observed particularly between *ZFP36, NFKBIA*, and *DUSP1*, key regulators of immune response signalling. In differentially expressed genes (bottom right), ranked by z-score, many key controllers of T-cell activation are upregulated in COVID-19, and the proliferation regulator *TXNIP* is downregulated.

## 6 Discussion

The MI estimation method we present here yields estimates that closely match analytical expectations, with accuracy approaching that of more computationally intensive kNN methods. For the case of the zero-inflated negative-binomial distribution, often observed in single-cell sequencing data, our method achieves parity with the kNN method. This enables precise quantification of non-linear regulatory relationships at the scale of modern single-cell sequencing capability.

To fully utilize the power of the approach, the data must be concentrated in low integer counts that follow (zero-inflated) negative binomial marginal distributions, which is an assumption that is violated when applying some processing techniques that involve re-scaling the data. However, whilst this violates the sampling assumptions of the NB distribution, the MI estimation procedure can be performed instead with empirical marginal distributions, leading to a loss in accuracy but with similar execution speed.

Furthermore, the minimum content per bin must be specified in the adaptive binning procedure, where too coarse a binning results in a biased estimate of the MI, and too fine a binning results in an estimate with high variance. However, as the binning is adaptive, this describes local smoothing rather than affecting the data distribution directly. For the experiments in this paper we have set this to be 50 and the estimates are consistent with expectations. There is further evidence that this, and variations in the estimated MI values overall, make little difference for regulatory network inference on bulk transcriptomics [7], however large modern single-cell datasets are potentially more sensitive to mis-estimation [27].

An advantage of a copula-based approach is that a *χ*^2^ test-statistic is available for minimal extra computation, where we compute the effective degrees of freedom for this test using a null joint uniform distribution. This enables hypothesis testing of all potential pairwise interactions and thus the rejection of spurious associations. As an example of how this can be applied, we study gene rewiring in naive CD4+ T cells in a SARS-CoV-2 single-cell sequencing study, calculating MI and *χ*^2^ test statistics for 2000 *×* 2000 highly-variable gene pairs in 77, 000 cells. This identified *ZFP36* as a key rewired gene between COVID-19 and healthy cells, with an environment that represents a program of tight regulation of T-cell activation.

## 8 Data availability

The SARS-CoV-2 dataset is available from its original source [17]. The algorithms described here for fast MI calculation can be found at github.com/dpohanlon/mimiq and on PyPI as `mimiq`, and for negative-binomial evaluation and fitting these can be found at github.com/dpohanlon/fast-nb and on PyPI as `fast-negative-binomial`. The code that can be used to produce the analysis and plots presented in the manuscript can be found at github.com/dpohanlon/mimiq-paper.

## A MI of zero-inflated variables

Two correlated random variables *X* and *Y* can be zero-inflated to generate the new variables

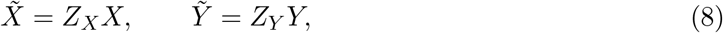

where *Z*_*X*_, *Z*_*Y*_ ∈ {0, 1} are Bernoulli variables with probabilities

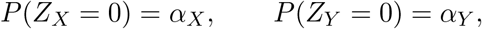

controlling the zero-inflation and independent of each other and of *X, Y*.

By definition the MI is

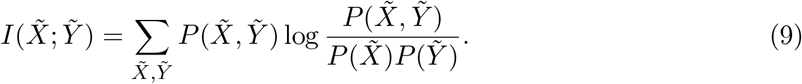

The sum can be separated into contributions where at least one of the variables is zero and those where both are positive,

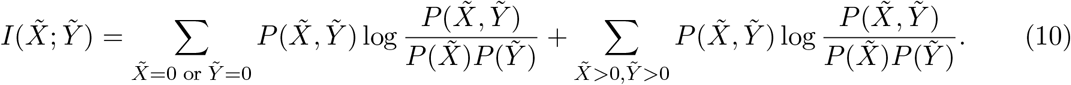

If the core distributions of *X* and *Y* take positive values with high probability,

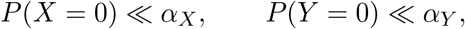

then terms in the first sum correspond predominantly to realizations where one of the Bernoulli variables is zero. Therefore they do not contribute to 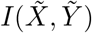 since *Z*_*X*_ and *Z*_*Y*_ are independent i.e. in this region 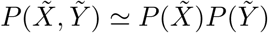 cancelling the logarithm in the first term of the sum.

The dominant contribution therefore comes from the region 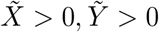, where necessarily *Z*_*X*_ = *Z*_*Y*_ = 1 and thus 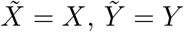. In this region

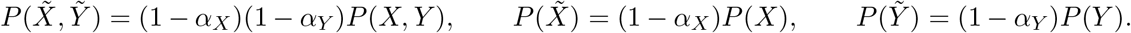

Substituting into the MI expression gives

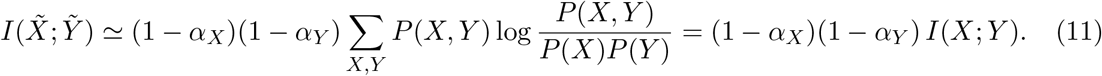

If the core variables *X* and *Y* are coupled through a Gaussian copula with correlation parameter *ρ*, then their MI is 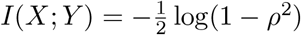. This gives the MI of the zero-inflated variables as

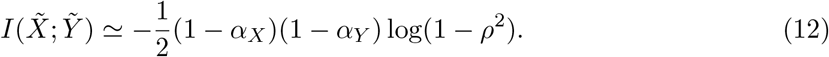

